# Distinctive epigenomic alterations in NF1-deficient cutaneous and plexiform neurofibromas drive differential MKK/P38 signaling

**DOI:** 10.1101/833467

**Authors:** Jamie L. Grit, Benjamin K. Johnson, Patrick S. Dischinger, Curt J. Essenburg, Stacy Campbell, Kai Pollard, Christine Pratilas, Tim J. Triche, Carrie R. Graveel, Matthew R. Steensma

**Author notes:** Corresponding Author, Van Andel Research Institute, 333 Bostwick Ave. NE, Grand Rapids, MI 49503. Contributed equally (Jamie L. Grit, Benjamin K. Johnson).

## Abstract

Benign peripheral nerve sheath tumors are the clinical hallmark of Neurofibromatosis Type 1. They account for substantial morbidity in NF1 and are difficult to manage. Cutaneous (CNF) and plexiform neurofibromas (PNF) share identical histology, but maintain different growth rates and risk of malignant conversion. The reasons for their disparate clinical behavior are not well explained on the basis of recent genome or transcriptome profiling studies. We hypothesized that CNFs and PNFs are epigenetically distinct tumor types that exhibit differential signaling due to genome-wide and site-specific methylation events. We interrogated the methylation profiles of 45 CNFs and 17 PNFs (Illumina EPIC 850K) using normal tissue controls from NF1 subjects. Based on these profiles, we confirm that CNFs and PNFs are epigenetically distinct tumors with broad differences in higher order chromatin states, and specific methylation events altering genes involved in key biological and cellular processes such as inflammatory mediator regulation of TRP channels, RAS/MAPK signaling, actin cytoskeleton rearrangement, and oxytocin signaling. Based our identification of 2 separate DMRs associated with alternative leading exons in *MAP2K3*, we demonstrate differential RAS/MKK3/P38 signaling between CNFs and PNFs. Epigenetic reinforcement of RAS/MKK/P38 was a defining characteristic of CNFs leading to pro-inflammatory signaling and chromatin conformational changes, whereas PNFs signaled predominantly through RAS/ERK. Tumor size also correlated with specific CpG methylation events. Taken together, these findings confirm that epigenetic regulation of RAS signaling fates accounts for observed differences in CNF and PNF clinical behavior. CNFs may also respond differently than PNFs to RAS-targeted therapeutics raising the possibility of targeting P38-mediated inflammation for CNF treatment.

## INTRODUCTION

Approximately 1 in 3,000 live births are affected by the tumor predisposition disorder, Neurofibromatosis Type 1 (NF1) making it the most common single gene-inherited disorder in humans. One of the clinical hallmarks and diagnostic *sine qua non* of NF1 is the formation of benign peripheral nerve tumors called, neurofibromas. Neurofibromas are a major cause of disfigurement, pain and morbidity within and outside of NF1. NF1 patients develop two types of neurofibromas: dermal or cutaneous neurofibromas (CNFs) that develop in the skin, and plexiform neurofibromas (PNFs) that arise from nerves situated in deeper anatomic compartments. Even though CNFs and PNFs share identical histology, they are pathophysiologically distinct. NF1-related PNFs are associated with the development of high grade malignant peripheral nerve sheath tumors, which are the leading cause of death of NF1 patients(1). In contrast, CNFs exhibit slower growth rates and are not associated with malignant potential(2-4). Although CNFs do not become malignant these benign tumors are often painful and disfiguring. These critical distinctions between CNFs and PNFs are not explained by genomic and transcriptomic studies that failed to identify consistent alterations in these tumors(5).

Previous genomic studies observed that approximately one third of CNFs exhibit focal chromosomal imbalance with a diversity of copy number alterations affecting transcription; however these findings were inconsistent among tumor samples(6). Expression profiling studies have yielded mixed results as to whether CNFs and PNFs can be discriminated through transcriptional means(5, 7), granted these studies were not powered specifically to address differences between CNFs and PNFs but rather PNFs and the derived malignancy, malignant peripheral nerve sheath tumor (MPNST). Even though CNFs and PNFs arise in distinct anatomic sites, mouse models have demonstrated that CNFs and PNFs share a common cell of origin (i.e. skin-derived precursors or SKPs(8) or GAP43+PLP+ precursor cells(9). Currently, it is not well understood how tumor microenvironment impacts neurofibromas development(10), however inflammation from NF1 haploinsufficient mast cells is strongly linked to PNF development(11). Prior work examining epigenetic modifications in CNFs and PNFs has also been confounded by the lack of tissue- and patient-matched controls which are not always available in the context of NF1-deficiency. Thus, there is insufficient data to adequately explain why CNFs and PNFs exhibit such distinct clinical behavior.

We now show that CNFs and PNFs are epigenetically distinct tumors that are distinguishable based on both site-specific and chromosomal-wide methylation differences. This work stands in contrast to prior studies that failed to identify differential patterns of methylation at the NF1 locus(12) in peripheral nerve tumors, and were underpowered to identify signaling impacts based on observed genome-wide methylation differences(13). Our work confirms that broad and distinct patterns of methylation result in differential signaling between CNFs and PNFs, as well as in the regulation of tumor size. Specifically, two differentially methylated regions (DMRs) in the *MAP2K3* and an upstream regulatory site for *MAPK14* are significantly altered between CNFs and PNFs leading to increased MKK3 and P38 expression, respectively. This epigenetic reinforcement of MKK3 and P38 expression leads to activation of both P38 and ERK in a validation cohort of CNFs. The MKK3/P38 signaling axis is linked to inflammation and pain signaling, as well as chromatin conformational changes through SWI/SNF complex regulation. Taken together, our data confirms that epigenetic regulation of key RAS signaling genes results in disordered growth and inflammation in the context of NF1 deficiency. These findings also confirm the importance of epigenetic regulation in CNF initiation and progression, as well as a potentially druggable signaling axis in MKK3/P38/ERK.

## METHODS

### Trial Participants and sample collection

45 cutaneous neurofibromas, 17 plexiform neurofibromas, and 9 normal skin and nerve samples were collected from individuals with a confirmed diagnosis of Neurofibromatosis Type 1. These samples were collected prospectively under an approved Spectrum Health/Van Andel Research Institute IRB protocol (SH/VAI IRB#2014-295) (NCT02777775). Additional specimens were analyzed according to ethical standards and under a Johns Hopkins Hospital (JHH) institutional review board (IRB)-approved protocol (JH IRB # J1649, PI Pratilas). The JH NF1 biospecimen repository is supported by a grant from the Neurofibromatosis Therapeutic Acceleration Program (NTAP, n-tap.org), to C.A.P. Analysis by Sage Bionetworks is supported through the Neurofibromatosis Therapeutic Acceleration Program (NTAP, n-tap.org). Informed consent was obtained from all participants. Tumor samples were isolated by microdissection to remove adjacent normal tissue then snap frozen in liquid nitrogen and stored at −80°C. Quality parameters included assessment of percent content (>95%) and viability (>90% nuclear viability) by H/E staining. Biospecimen handling was performed according to BRISQ guidelines.

### 5mC interrogation by Infinium MethylationEPIC array

To extract DNA, frozen tissue was manually dissected into small pieces and homogenized by bead beating (Lysing Matrix D; MP Biomedicals) in UltraPure Phenol:Chloroform:Isoamyl Alcohol (ThermoFisher) according to the manufacture’s protocol. DNA was quantified by Qubit fluorometry (Life Technologies) and 500ng of DNA from each sample was bisulfite converted using the Zymo EZ DNA Methylation Kit (Zymo Research, Irvine, CA USA) following the manufacturer’s protocol using the specified modifications for the Illumina Infinium Methylation Assay. After conversion, all bisulfite reactions were cleaned using the Zymo-Spin binding columns, and eluted in 12 uL of Tris buffer. Following elution, BS converted DNA was processed through the EPIC array protocol. The EPIC array contains >850K probes querying methylation sites including CpG islands and non-island regions, RefSeq genes, ENCODE open chromatin, ENCODE transcription factor binding sites, and FANTOM5 enhancers. To perform the assay, 7uL of converted DNA was denatured with 1ul 0.4N sodium hydroxide. DNA was then amplified, hybridized to the EPIC bead chip, and an extension reaction was performed using flurophore labeled nucleotides per the manufacturer’s protocol. Array beadchips were scanned on the Illumina iScan platform and probe specific calls were made using Illumina Genome Studio software.

### Western Blotting

To select a subset of tumor samples for protein analysis, samples were divided into methylation high, methylation intermediate, and methylation low groups based on the beta values across MAP2K3 DMR1 and samples from each group were randomly chosen. Protein lysates were prepared by manually homogenizing frozen tissue in RIPA buffer containing protease and phosphatase inhibitor cocktails (Roche). Proteins were separated on a 4-20% TGX SDS-PAGE gel (Bio-Rad) and transferred to a PVDF membrane (Invitrogen). Blots were blocked in 5% dry milk in TBST buffer (20 mmol/L Tris-HCl pH 7.4, 150 mmol/L NaCl, 0.1% Tween-20) and incubated at 4°C overnight in primary antibody; MKK3 (Cell Signaling #5674), p38 (Cell Signaling #9219), phospho-p38 (Cell Signaling # 4511), phospho-ERK1/2 (Cell Signaling #9101), and β-Actin (Cell Signaling #3700). Densitometry analysis was performed in ImageJ.

### EPIC methylation array data pre-processing

Data were analyzed using a modified workflow that is similar to the ChAMP methylation array analysis procedure in R (v3.5.1). Briefly, samples were filtered for probes with poor or skewed intensities (detection P-value < 0.01) and entire samples were removed from the dataset if they contained >10% failed probes. One sample exceeded the aforementioned filtering criteria and was removed from the dataset resulting in 70 total samples. Next, probes that have previously been identified to skew downstream differential methylation analyses (SNP probes, cross-reactive probes with other genomic regions, etc.) were removed in addition to probes that target sex chromosomes. A batch effect was detected based on SVD analysis and corrected using the sva (v3.30.1) package in R. In total, 717,148 probes were analyzed for differential methylation across tissue types.

### Differential methylation analysis

Differentially methylated loci between cutaneous and plexiform neurofibromas were identified using a hierarchical generalized linear mixed model (GLMM) approach with a logit link function as implemented in glmmTMB (v0.2.2) on the pre-processed beta values. Partially repeated tissue sampling was modeled as a random effect with patient nested within methylation array slide unless specified otherwise. Group-level differences were determined using a likelihood ratio test (LRT) with a significance threshold of q < 0.05. False discovery rate adjustment was done using the Benjamini-Hochberg procedure. Models were filtered out for downstream analysis if they failed to converge. In total, 31,201 probes were found to be differentially methylated between cutaneous and plexiform neurofibromas. Differentially methylated regions were called using DMRcate (v1.18.0). Results from glmmTMB were wrangled into a suitable data structure as input for DMRcate by using the Wald statistic as the stat and a quasi-beta fold-change using the exponentiated model estimates for each probe. Default parameters were used for DMRcate with the bandwidth scaling factor (C parameter) set to 2.

To identify differentially methylated loci associated with cutaneous neurofibroma size, a GLMM with a logit link was fitted, adjusting for age and sex differences and a random intercept term for partially repeated tissue sampling. Significant associations (q < 0.05) between CNF size in millimeters and probe-level methylation were determined as described above. A total of 188 loci were found to be significant. Positive or negative correlations were computed on significant probes using Kendall’s Tau. False discovery rate was controlled using the Benamini-Hochberg procedure and significant correlations were determined at a q < 0.05 threshold. Significantly correlated probes were filtered on a delta Beta-value (maximum Beta-value minus the minimum Beta-value) of 0.2, resulting in 34 loci.

### Inference of chromatin conformation from EPIC methylation arrays

Chromatin compartments were computed at 100kb resolution as previously described and implemented in compartmap (v1.65.71). Briefly, pre-processed M-values were subset to “open sea” CpG probes (at least 4kb away from annotated CpG island) and masked probes that were found in at least 50% of samples were imputed using k-nearest neighbor via the impute (v1.59.0) R package. Next, loci were median summarized in 100kb bins. Group-level compartments were inferred by computing Pearson correlations of summarized bins and the first principal component of the correlation matrix. A/B compartments correspond to positive (open chromatin) and negative (closed chromatin) eigenvalues, respectively. Genome-wide discordant compartments were identified by comparing the sign of the eigenvalue for overlapping genomic bins and filtering out those with small absolute eigenvalues (>0.02). In total, we identified 2937 discordant chromatin compartments between plexiform and cutaneous neurofibromas. Results were plotted using circlize (v0.4.8).

### Pathway and GO term enrichment

Enrichment of pathways and gene ontology (GO) terms was performed using the gometh function within the missMethyl (v1.16.0) package in R. Briefly, gometh considers the relatively uneven density of loci covered on the Infinium methylation arrays and utilizes this information when computing enrichment, similar to the approach goseq uses for RNA-seq. Significant CpGs were identified as described above and the background probe set was derived following pre-processing. Significant GO terms and KEGG pathways were determined using a q < 0.05 threshold. Results were plotted using ggplot2 (v3.2.1).

### Tissue purity estimates

Tissue purity was estimated by PAMES (v0.2.3) and annotations built for computing informative sites using IlluminaHumanMethylationEPICanno.ilm10b2.hg19 (v0.6.0). Cutaneous neurofibromas were compared against normal skin samples and plexiform neurofibromas were compared against normal nerve. Results were plotted using ggplot2 (v3.2.1).

### Copy number estimation from EPIC methylation arrays

Copy number alterations were computed using SeSAMe (v1.3.2) with minor modifications for plotting functionality. Briefly, data were processed using the “open SeSAMe” procedure, producing a signal set object. Next, samples were segmented and called for copy number differences, comparing CNFs to normal skin and PNFs to normal nerve. The genome annotation used was hg19.

### RNA-sequencing data analysis

Paired-end, raw neurofibroma and plexiform neurofibroma RNA-seq data were downloaded from syn4939902. Sequencing lanes were merged, followed by alignment with STAR (v2.7.0f) to b37 (downloaded from the GATK resource bundle-https://software.broadinstitute.org/gatk/download/bundle), using the Gencode v19 annotations. Alignment was performed using default parameters with the following modifications: --twopassMode Basic, --outSAMtype BAM SortedByCoordinate, and --quantMode GeneCounts. Reverse-stranded gene counts were read into R (v3.6.1) using edgeR (v3.27.14), excluding sample 2-025 due to sample quality. Gene counts were restricted to known, protein coding genes and lincRNA. Samples were further filtered to genes that had greater than 1 count per million (CPM) in at least 3 samples. Libraries were normalized using the trimmed mean of M-values method. Dimensionality reduction was performed using the prcomp function (v3.6.1) on the filtered log2 CPM and plotted using ggplot2 (v3.2.1).

### Data availability

Raw and processed EPIC array data are available from Synapse: syn4939910. Raw RNA-seq fastq data were downloaded from syn4939902.

### Statistical Methods

Protein expression and methylation correlations were done using R version 3.5.1.

## RESULTS

### CNFs and PNFs have distinct global methylation profiles

CNFs are confined to the skin and are a hallmark of NF1. These benign tumors typically arise in puberty and vary dramatically, in which NF1 patients can have between one or thousands of CNFs that cover the body. Even though the size of CNFs can vary in NF1 patients (Fig 1a), CNFs are associated with limited growth (<3 cm). The only effective treatment option for CNFs is surgical removal, however this is impracticable in patients with severe tumor burden. CNFs are composed of neoplastic Schwann cells, mast cells, and fibroblasts (Figure 1a, middle and lower panels). In addition, CNFs often include a collagenous and myxoid extracellular matrix. CNFs and PNFs cannot be differentiated histologically and require clinical context to make an accurate diagnosis of a surgical sample(14). PNF growth is dysregulated, can develop anywhere in the body, and can extend along nerve branches. Because of these features, PNFs can compress nearby structures and cause pain and disfigurement. For example, a PNF included in this study was present in an axial location where it was growing in an unchecked manner (Fig 1b). The risk of CNF-associated malignancy is extremely low, whereas PNFs are able to dedifferentiate into aggressive sarcomas in 8-13% of patients(15).

**Figure 1.**
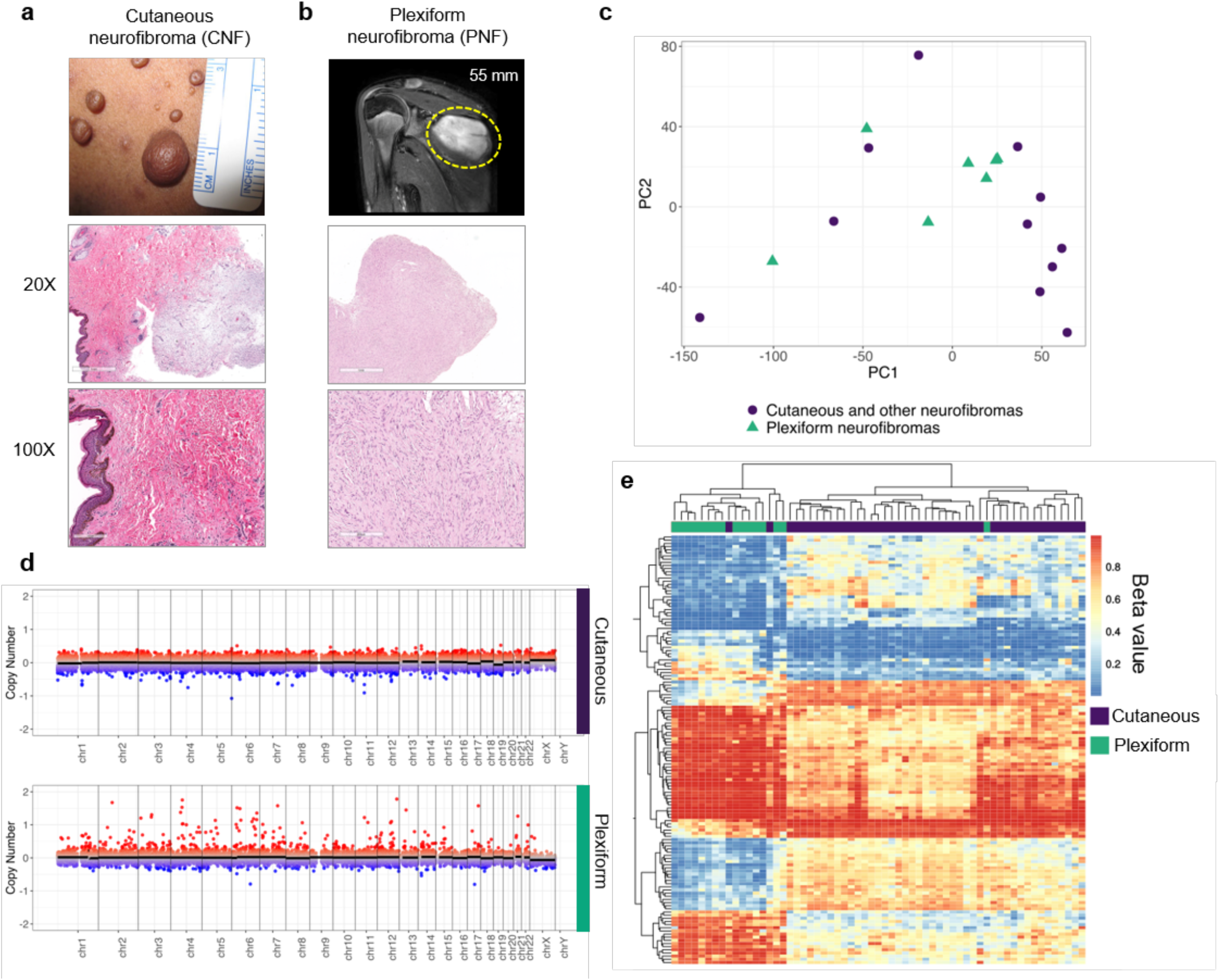

In order to explain these pathophysiological differences, we compared publicly available RNA-seq of a test cohort of cutaneous and other NF1-deficient non-plexiform neurofibromas and PNFs. Principal Component Analysis (PCA) of the RNA-seq data confirms that histologically identical cutaneous and other NF1 deficient non-plexiform neurofibromas and PNFs cannot be discriminated on the basis of global gene expression profiles largely due to transcript heterogeneity (Fig 1c). Prior expression profiling studies using cDNA microarrays failed to identify distinct CNF and PNF transcriptome signatures at the macro level, even though individual differences in key gene expression were identified(5). We also evaluated whether genomic differences were present among our CNF and PNF cohorts and did not observe any large-scale differences in copy number variation (Fig 1d). These results confirm that few distinctive genomic or transcriptome alterations exist outside of putative NF1 deficiency (Fig 1c-d).

DNA methylation changes play key roles in development, disease and aging, with the added benefit of being a highly stable epigenetic mark that can be readily assayed, making this a clinically actionable and functional readout(16-20). Thus, we determined the methylation profiles using the Infinium MethylationEPIC array in a discovery cohort of 62 CNF and PNF samples from a total of 38 patients. The MethylationEPIC array is a targeted approach to interrogate approximately 850,000 methylation sites throughout the genome, including coding and regulatory space (e.g. FANTOM5 annotated enhancers, etc.) in addition to CpG islands, shores, and shelves. We applied a hierarchical generalized linear mixed effects model to identify differentially methylated loci between CNFs and PNFs, controlling for age and sex differences with a nested random effect to control for partially repeated measures. In total, we identified 31,201 significant differentially methylated probes (DMP; q < 0.05). By examining differentially methylated loci with an absolute odds ratio greater than 4 and clustering using a semi-supervised hierarchical approach, we establish a distinct, base-pair resolution epigenetic signature for CNF and PNFs (Fig 1e). This probe-based analysis confirms that individual CpG-based methylation events are highly consistent within tumor types, yet between CNFs and PNFs there are clearly definable, distinct global methylation profiles.

### CNFs and PNFs display distinctive 3D chromatin architecture

Next, we sought to determine if nuclear organization and higher-ordered chromatin differed between tumor types. Chromosomal DNA is organized into A/B compartments that largely correspond to being either transcriptionally active DNA compartments (A compartment) or silenced DNA compartments (B compartment)(21). As such, DNA compartmentalization is a critical determinant of gene expression leading to cell fate determination, and organized tissue development(22, 23). Typically, chromatin compartments are identified using HiC or other assays to directly measure long-range chromatin contacts(24). However, it has been shown that Infinium Human Methylation 450k or EPIC array can reconstruct higher-order chromatin structure similar to HiC(25). Thus, we inferred A/B group-level compartments in CNF and PNFs at 100kb resolution. We show that the 3D genomic organization between CNFs and PNFs is 80-85% concordant. However, we determined that approximately 15-20% of the compartments are inverted or discordant between CNF and PNFs. This discordance of PNF and CNFs compartments is observed genome-wide (X chromosomes not assessed) (Fig 2a). These data provide additional support that CNFs and PNFs possess distinct epigenomes. Given that individual CpG methylation changes and associated effects are difficult to interpret in isolation unless they are placed in the context of neighboring CpG loci and summarized into region-level changes, we used the differentially methylated probes identified above to call 6,097 significantly differentially methylated regions (DMRs; q < 0.05) using DMRcate. A focused evaluation of the top 250 DMRs in terms of fold change revealed that 100 sites were strongly associated with gene regulation (i.e promoter/enhancer space). We observed that areas of promoter/enhancer overlap were evenly distributed across all chromosomes except chromosome 17 which was disproportionately affected (Fig 2cd). Discordant regulatory DMRs spanned the entire length of Chromosome 17, including key genes such as *NF1, TP53*, and *MAP2K3* (Fig 2d). Taking into account all DMRs, chromatin structural analysis reveals a broad distribution that extended well beyond the transcriptional start site, including CpG islands, shores, and shelves (Fig 2e). Taken together, we provide evidence that associates NF1-deficiency in otherwise transcriptionally and genomically similar lesion types, can exhibit altered 3D genomic organization and higher order chromatin structure. These results imply that NF1-deficiency impacts signaling events through methylation changes that can directly remodel the chromatin architecture throughout the genome. This global impact of NF1-deficiency may underlie the morphological and pathological variation of CNFs and PNFs in NF1 patients.

**Figure 2.**
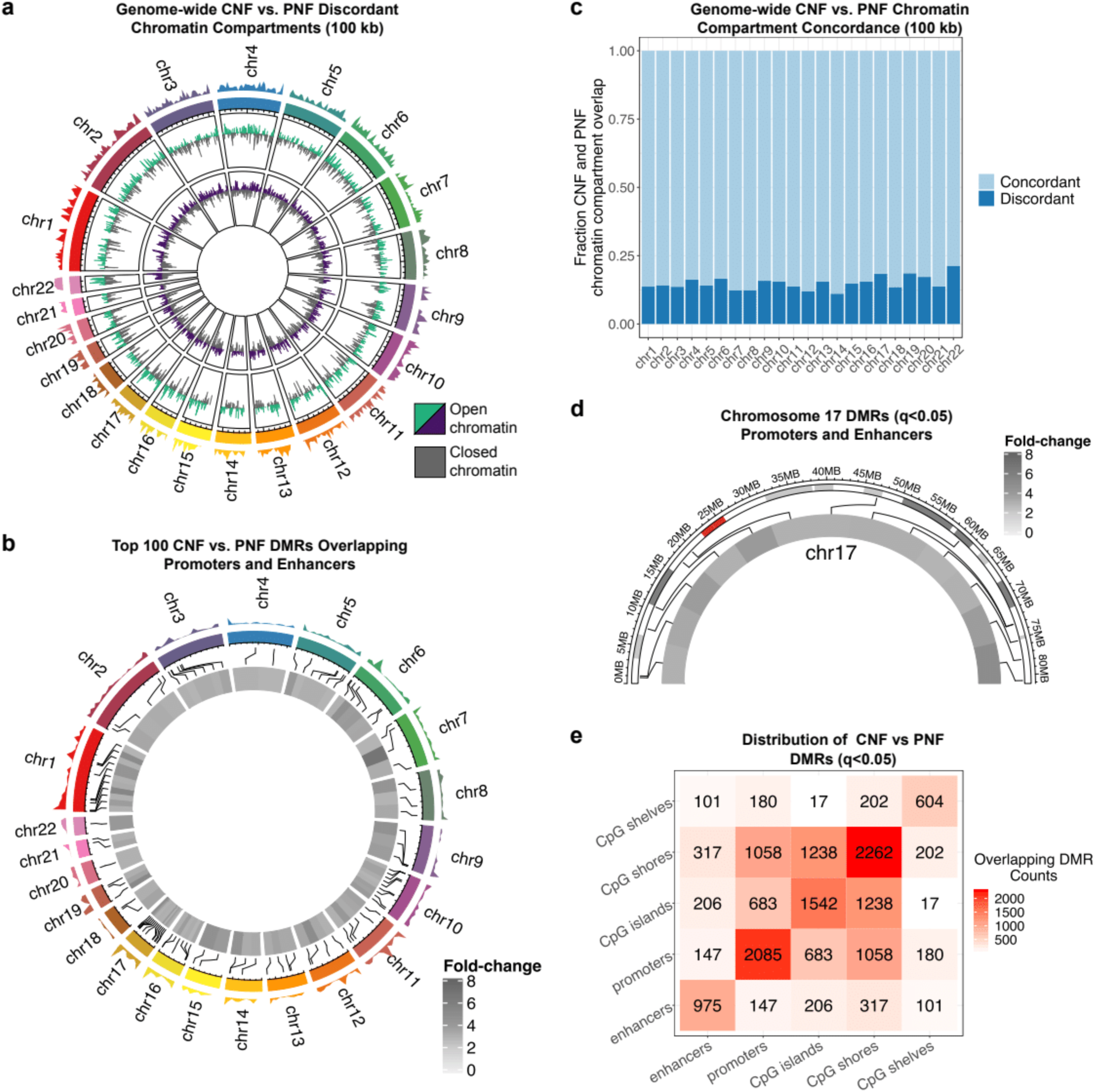

### CNF size variation is correlated with differential methylation

The CNF size variation within and between NF1 patients is striking and can even be discordant among monozygotic twins(26). To determine if CNF size correlates with site-specific methylation events, we collected CNFs of various sizes. Adjusting for age and sex differences, we determined significant associations (q < 0.05) between CNF size in millimeters and probe-level methylation status. A total of 188 loci were found to be significant. Both positive and negative correlations were discovered among the 188 associated loci, with 34 achieving an effect size that also reached statistical significance (Fig 3). These results indicate that CNF size is significantly influenced by methylation of specific loci, which sheds light on the significant variance observed clinically.

**Figure 3.**
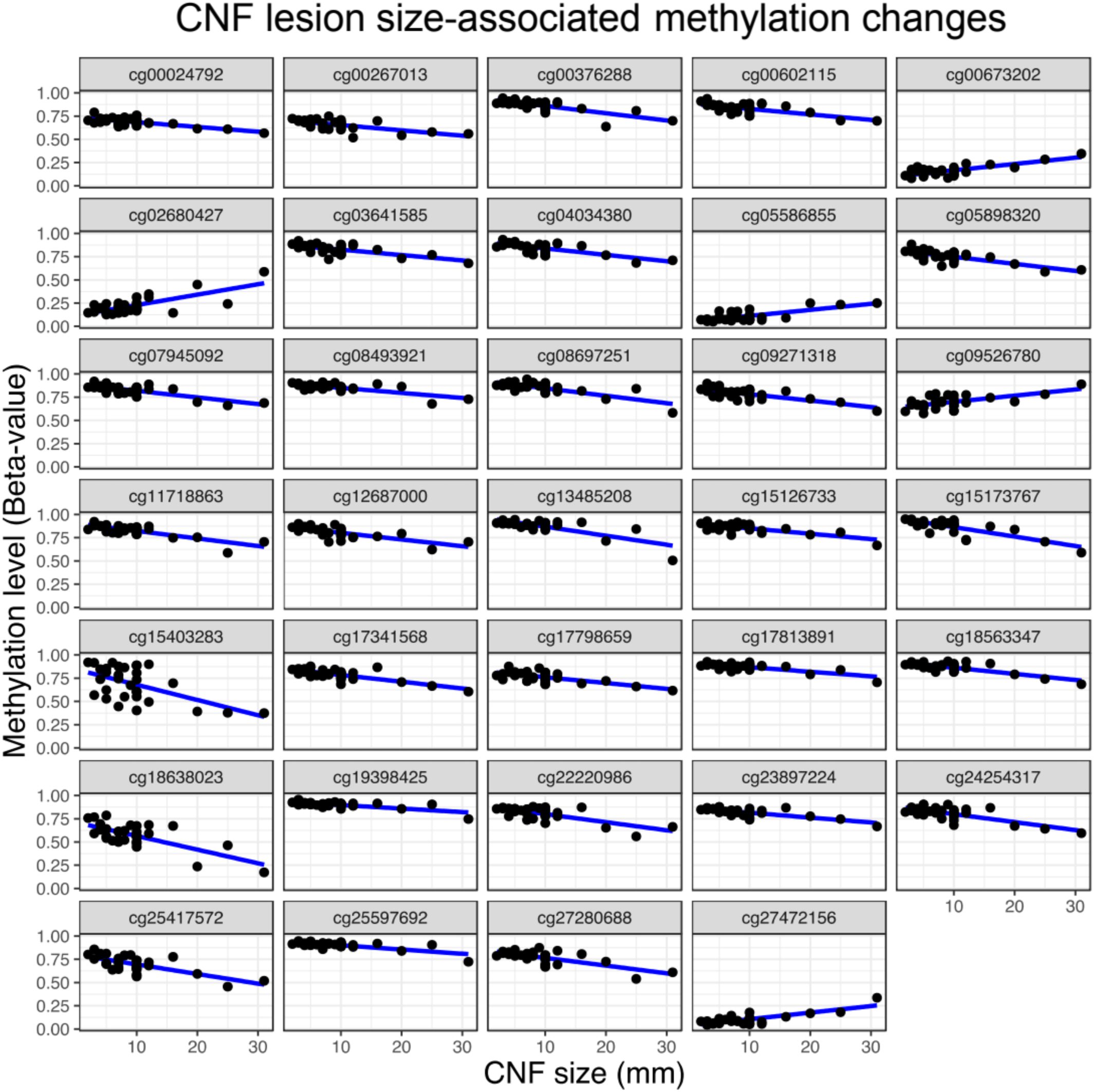

### Differentially methylated loci are enriched in inflammatory and pain signaling pathways

To understand how these methylation patterns may promote CNF and PNF development through cell signaling alterations, we examined CNF- or PNF-specific DMR patterns associations with specific pathways. Kyoto Encyclopedia of Genes and Genomes (KEGG) pathway enrichment analysis is a high-level functional analysis of epigenomic data that allows for annotation of key cell signaling and biological processes. KEGG analysis of global DMR data identified several key cellular processes that differed significantly between tumor types as a result of predicted expression impacts from methylation events. CNFs and PNFs demonstrate differential regulation of inflammatory mediators of transient receptor potential (TRP) channels, RAS-mediated growth and proliferation, actin cytoskeleton, and somewhat unexpectedly, oxytocin signaling (Fig 4a). To functionalize these results and validate a role for individual DNA methylation events in tumor-type associated inflammation and pain signaling, we examined significantly correlated DMRs associated with genes in the Inflammatory Mediator Regulation of TRP Channels KEGG pathway. Two highly significant DMRs (DMR1 p = 2.72E-21; DMR2 p = 1.84E-08) were discovered within the primary and alternative promotors and leading exons of the *MAP2K3* gene that encodes the MAP Kinase Kinase, MKK3 (Fig 4b). Across DMR1, PNFs displayed higher DNA methylation, while CNFs were more highly methylated in DMR2 (Fig 4c). As MKK3 plays a role in pain signaling, inflammation and cancer(27, 28), we hypothesized that methylation of DMR1 and DMR 2 would instruct MKK3 protein expression leading to differential RAS signaling. Western blot of a representative subset of CNF and PNF tumors demonstrated that MKK3 expression was highly correlated to *MAP2K3* DMR methylation status (DMR1:MKK3 ρ = −0.85, p = 0.00018; DMR2:MKK3 ρ = 0.68, p = 0.0084) (Fig 4d). This strong inverse correlation points to exquisite epigenetic control of MKK3 expression through DMR events occurring at sites of alternative leading exons.

**Figure 4.**
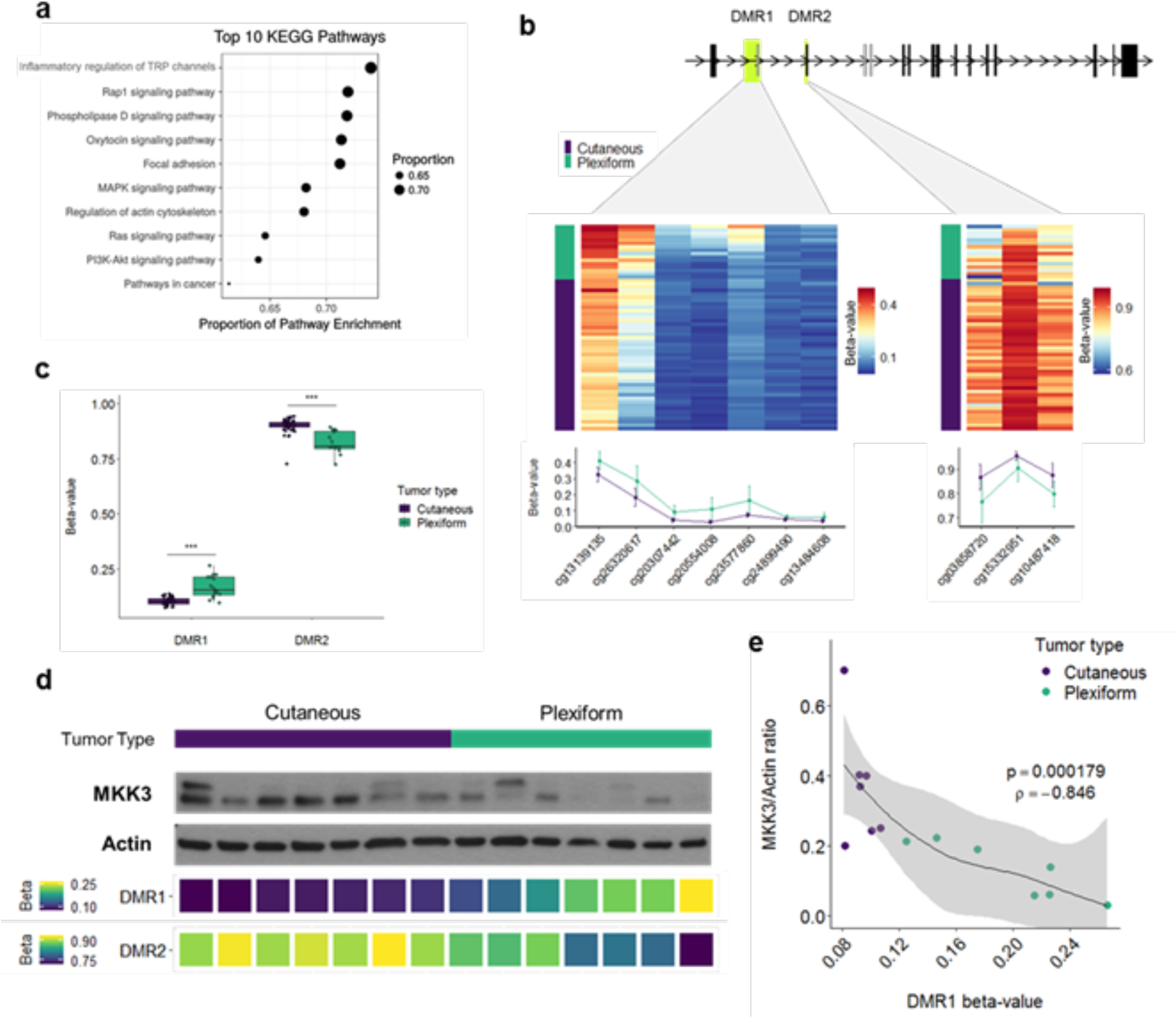

As p38 is the primary effector of MKK3 signaling in response to cellular stress and cytokine stimulation(29), as well as chromatin conformational changes such as those observed in Figure 2, we hypothesized that DNA methylation also regulates p38 expression in CNFs and PNFs. Because methylation events can also impact kinase activation(30), we also assessed whether *MAP2K3* methylation events affected downstream P38 activation. We identified a significant DMR (p = 1.66E-16) approximately 3.5 kb upstream of the *MAPK14* gene, which encodes the p38α isoform. Although the DMR lies within the promotor region of SLC26A8 gene, p38 protein levels were negatively correlated with methylation at this site (r = −0.56, p = 0.038). Methylation was higher in the PNF group than the CNF group leading to variable but decreased P38 expression (T180/T182) in PNFs (Fig 5a). Phospho-P38 expression positively correlated with methylation of the *MAP2K3* DMR1 suggesting that DNA methylation regulates both expression and activation of p38 in CNFs and PNFs. More specifically, CNFs were observed to overexpress both MKK3 and P38 leading to consistent P38 activation. Moreover, P38 activation closely correlated with pERK expression suggesting possible cross talk between RAS/ERK and MKK3/P38.

**Figure 5.**
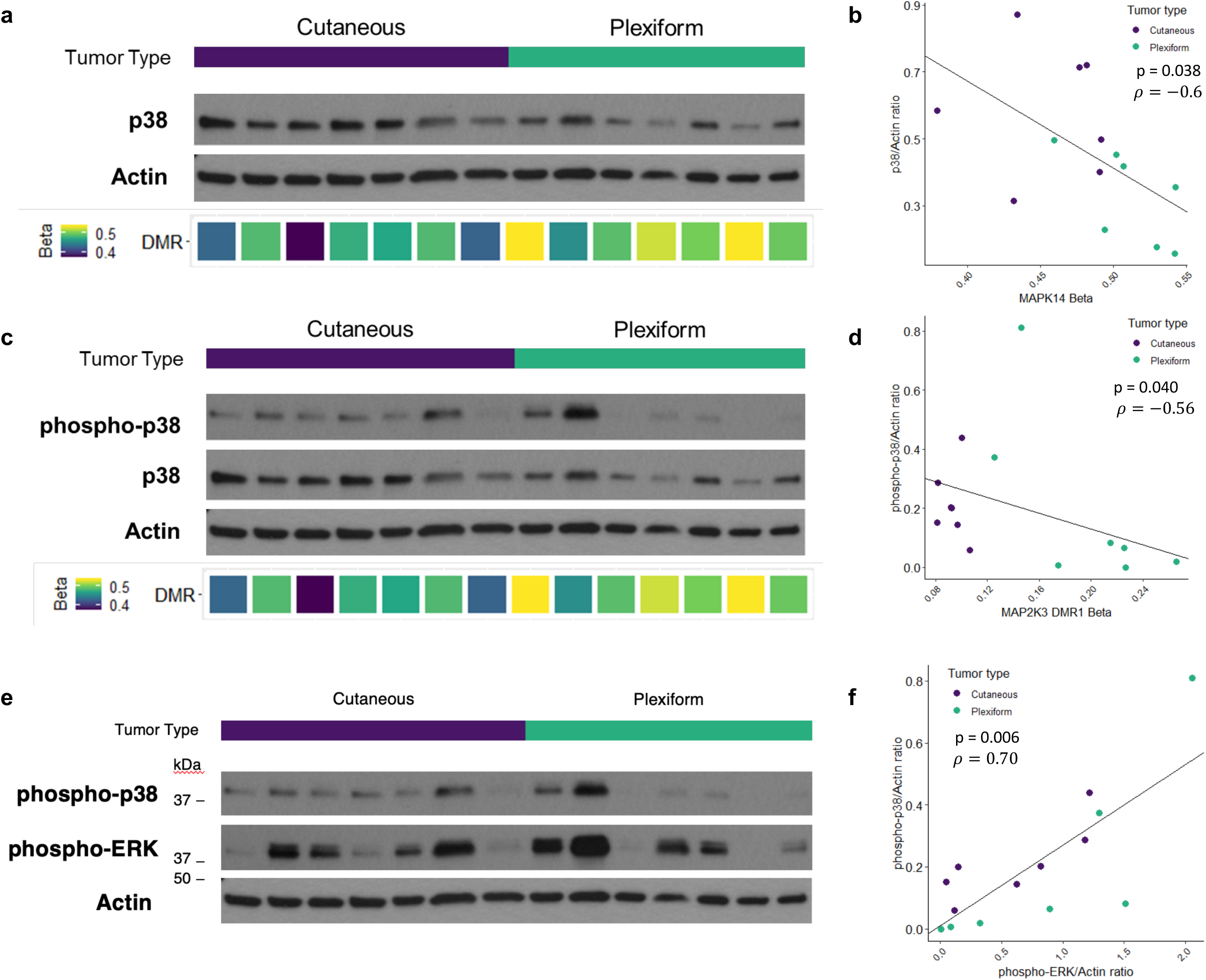

## DISCUSSION

This study represents the largest epigenetic analysis of CNFs and PNFs to date. Greater than 99% of NF1 patients exhibit both CNFs and PNFs over the course of their lifetime accounting for a substantial negative impact on quality of life(4). Pain is a constant feature of both neurofibroma subtypes, yet how pain signaling occurs in peripheral nerve tumors is poorly understood. Despite the recent demonstration of MEK inhibitor effectiveness in PNF treatment(31), it is unclear whether CNFs respond with equal efficacy. More therapies are needed to treat neurofibroma tumor progression and pain as both of these clinical features contribute significantly to morbidity in NF1. Our findings address an unmet clinical need for neurofibroma treatment and offer significant mechanistic insight into how benign nerve tumors initiate, progress, and generate symptoms through epigenetic means.

In the absence of consistently identifiable transcriptomic or genomic alterations in CNFs and PNFs, our work confirms that methylation events are key molecular determinants of nerve tumor initiation, growth, and pain generation. How epigenetic regulation of kinase signaling affects cancer predisposition in these tumors remains unclear, however recent data confirms that accumulating epigenetic alterations in a single field or region are associated with elevated cancer risk(32-36). These data lay the groundwork for future studies examining how epigenetic alterations affect PNF conversion into MPNSTs, as well as protective mechanisms that spare CNFs from cancerous progression or unchecked tumor growth. The identification of robust differences in the methylation profiles of CNFs versus PNFs further confirms their distinct biology, laying the groundwork for future development of clinical biomarkers.

Chromatin conformational states differed significantly between CNFs and PNFs and were strongly linked to both site-specific and geographic-specific methylation events. These findings suggest that chromatin accessibility may broadly affect gene expression in CNFs and PNFs. More work is needed to determine how epigenetic alterations affect regulatory genes that are known to contribute to tumor size and ultimately, cancer predisposition. Based on our probe-based analysis of tumor tissue, we identified 34 CpG methylation sites that were statistically correlated with CNF size. Unfortunately, the genes corresponding to the individual methylation sites could not be identified with statistical confidence, nor could we link these methylation events with specific biological processes or signaling pathways. Regardless, these data confirm that CpG methylation influences CNF tumor size, possibly through a novel mechanism. More work is needed in this area.

Our data confirms that CNFs and PNFs strongly exhibit differential methylation at two established DMR’s (i.e. DMR1 and DMR2) that are situated immediately upstream of the *MAP2K3* transcriptional start site. This pattern of differential methylation resulted in upregulated expression of MKK3 and P38 in CNFs, whereas in PNFs the reciprocal effect was observed with downregulated expression (Fig 5). This effect was consistent within and across tumor types despite expected signaling heterogeneity from analyzing whole tumor tissue. The cell types that contributed to the observed differences in methylation profiles could not be determined. Unfortunately, deconvolution analysis is dependent on cell type-specific profiles which are lacking for neurofibromas.

*MAP2K3* was previously identified as a candidate imprinted gene in the context of NF1 deficiency(37), but the roles of its upstream DMRs is not well characterized. DMR1 is generally thought to regulate expression of an alternative coding region with sequence homology to exon 1, whereas DMR 2 regulates exon 1 directly. The importance of alternative exon expression in cancer is increasingly being recognized as it has been used to identify breast cancer subtypes using RNA seq data from the The Cancer Genome Atlas (TCGA) Breast Invasive Carcinoma (BRCA) cohort(37). DNA methylation status was also shown to affect expression of alternative exons in the sphingosine 1-phosphate (SPHK1) gene in gastroesophageal cancer(38, 39). Apart from these studies, the impact of alternative exon expression on tumorigenesis has not been well described.

Our work extends these important findings by identifying alternative exon utilization as a potential regulatory mechanism for the MKK3/P38 signaling axis. More broadly, these data strongly point towards epigenetic control of RAS signaling fates downstream of NF1. We propose a schema where P38 activation in response to cellular stress and cytokine signaling inputs is reinforced in CNFs, whereas PNFs appear to signal predominantly through RAS/MEK/ERK leading to growth and proliferation (Fig 6-schematic). Future studies are needed to better define the implications of P38 activation in neurofibromas and their various cellular constituents. It is important to note, however, that crosstalk between the MKK3/P38 and RAS/MAPK signaling pathways has not been extensively studied. Prior work suggests a potential inhibitory role for RAS/ERK in mitigating P38-mediated inflammation(40). Interestingly, P38 is not typically activated in response to mitogenic stimuli, but we observed a high degree of correlation between pP38 and pERK expression. These results suggest that differential methylation may enhance crosstalk between MKK3/P38 and RAS/ERK leading to mixed signaling effects.

**Figure 6.**
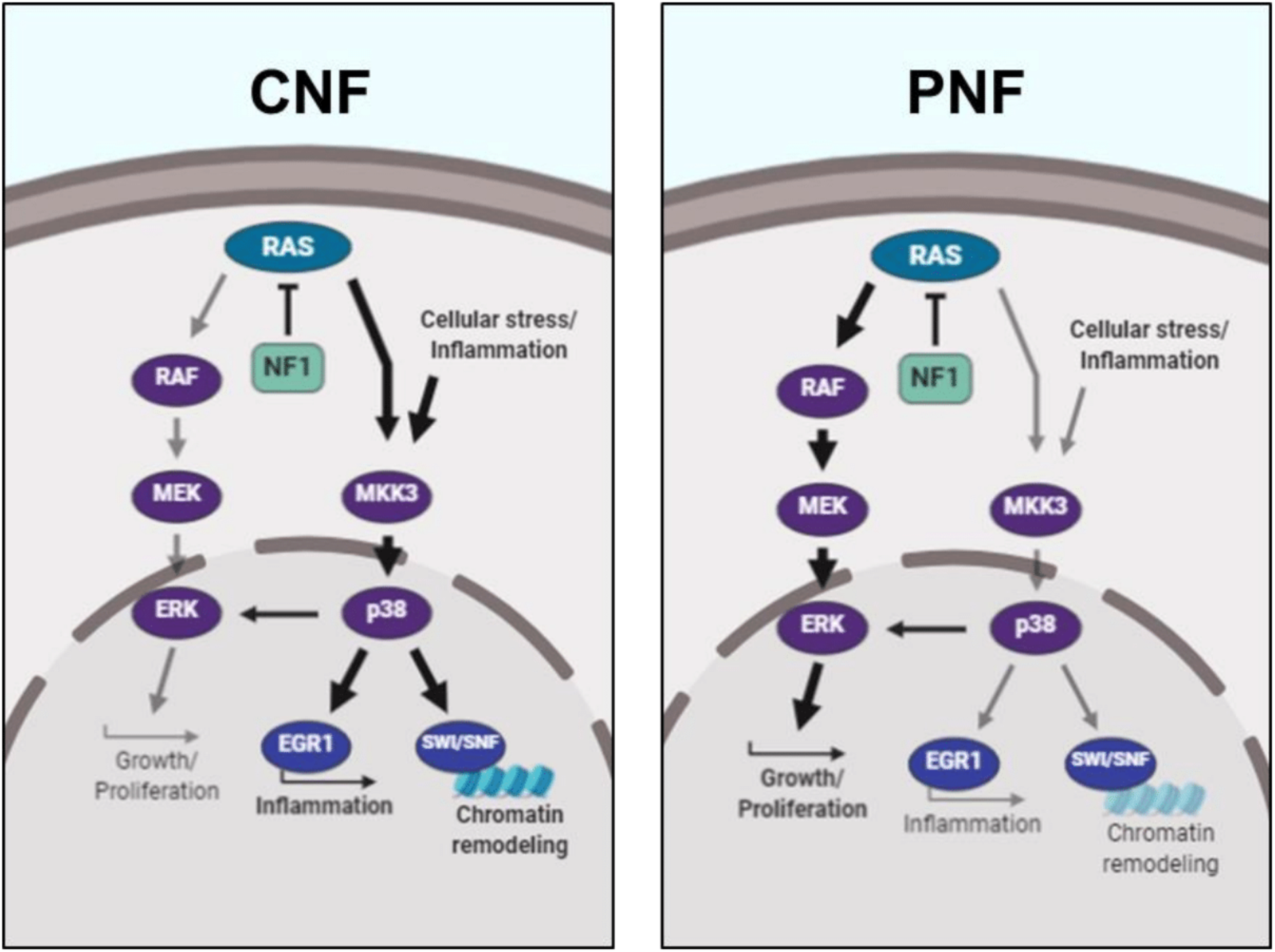

Proof of this concept comes from our observation that upregulated MKK3 expression, in turn, correlated with both P38 expression (P38) and activation (pP38) indicating strong epigenetic reinforcement of the MKK3/P38 axis in CNFs (Figure 4). Two expected results of P38 activation are activation of the MKK3/P38/EGR1 inflammatory cascade(41) and changes in chromatin conformation mediated through the SWI-SNF complex family(42). Relevant to the role of EGR1 in the pro-inflammatory response, it is intriguing that in our unbiased gene set enrichment analysis we identified inflammatory mediator regulation of TRP channels and phospholipase D signaling as the most significant altered signaling pathways related to DMRs (Figure 4a), granted EGR1, itself, was not found to be differentially methylated (data not shown). Pain is a constant feature of CNFs and PNFs leading to significant morbidity. Pain signaling in nerve tumors is not well understood and difficult to manage, clinically. These data identify a potentially novel mechanism for epigenetic regulation of pain signaling in nerve tumors and a targetable signaling axis in MKK3/P38.

P38 is involved in the direct recruitment of SWI/SNF complexes to gene promoters resulting in chromatin modification and enhanced expression (15208625). Although the methylation states of SWI/SNF complex family member DMRs were not discordant between CNFs and PNFs, it is plausible that reinforced MKK3/P38 signaling would exert its effect through SWI/SNF leading to the observed conformational changes. Further studies are needed to determine how SWI/SNF affects expression of genes involved in growth, proliferation, and inflammation. Moreover, the effects of targeting P38 may be amplified by expected loss of recruitment of SWI/SNF complexes to target genes.

## CONCLUSION

The epigenetic distinctions between CNFs and PNFs extend from the level of chromatin conformational change down to altered expression of genes that regulate or modulate RAS signaling. These findings are intriguing given that the analyzed tumors arose in the context of RAS deregulation as a result of NF1-deficiency. Based on KEGG pathway analysis, it is likely that methylation events are involved in regulation of pain signaling down to the level of inflammatory mediator production. More work is needed in many aspects of neurofibroma epigenetics, including studies targeting P38 and its downstream effectors. As such, we present a new signaling paradigm where differential methylation between tumor types results in reinforcement of inflammatory signaling in CNFs, and classical RAS/MEK/ERK activation towards growth in PNFs.

## Acknowledgements

The authors thank the Van Andel Genomics Core for providing facilities and services.

