## Supplemental for "Distinctive epigenomic alterations in NF1-deficient cutaneous and plexiform neurofibromas drive differential MKK/P38 signaling"

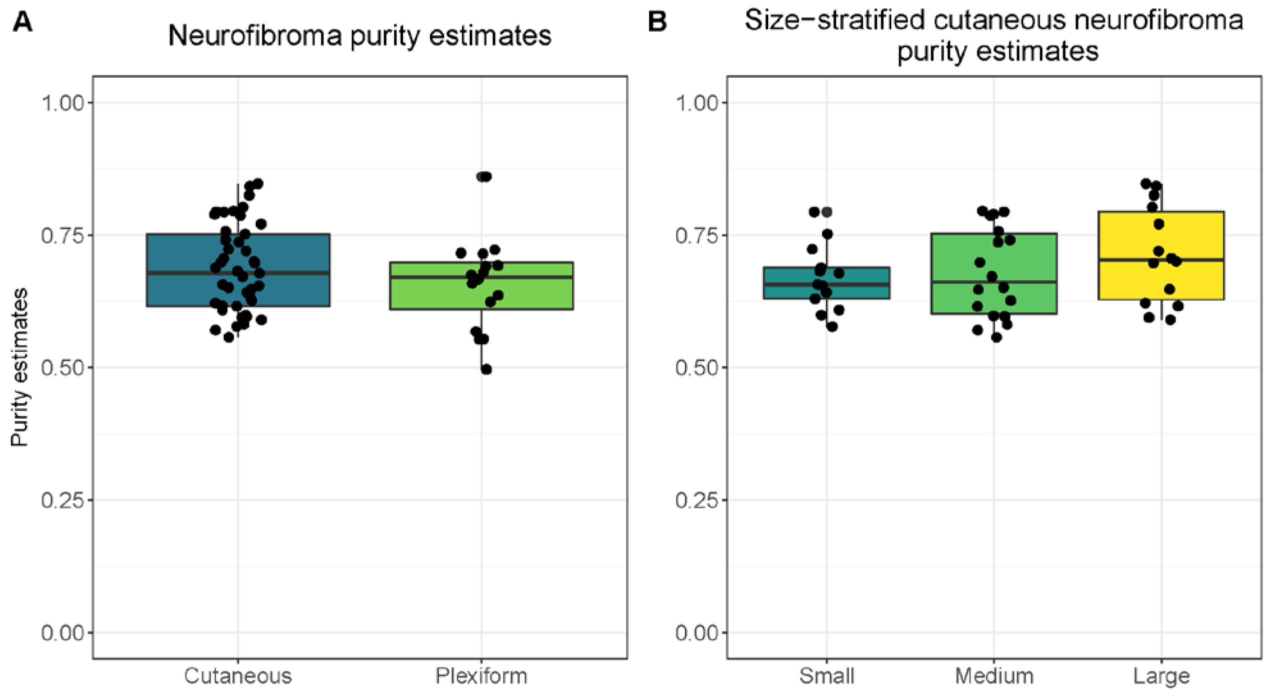


**Supplemental Figure 1: Purity analysis CNF and PNF tissue.** a) Cutaneous and plexiform neurofibroma methylation profiles are compared against normal controls. Purity estimates indicate significant enrichment of tumor tissue in queried samples. b) Size Comparison reveals equal purity among the three categories of tumors: incipient (<5mm), medium (5-10mm) and large (>10mm).

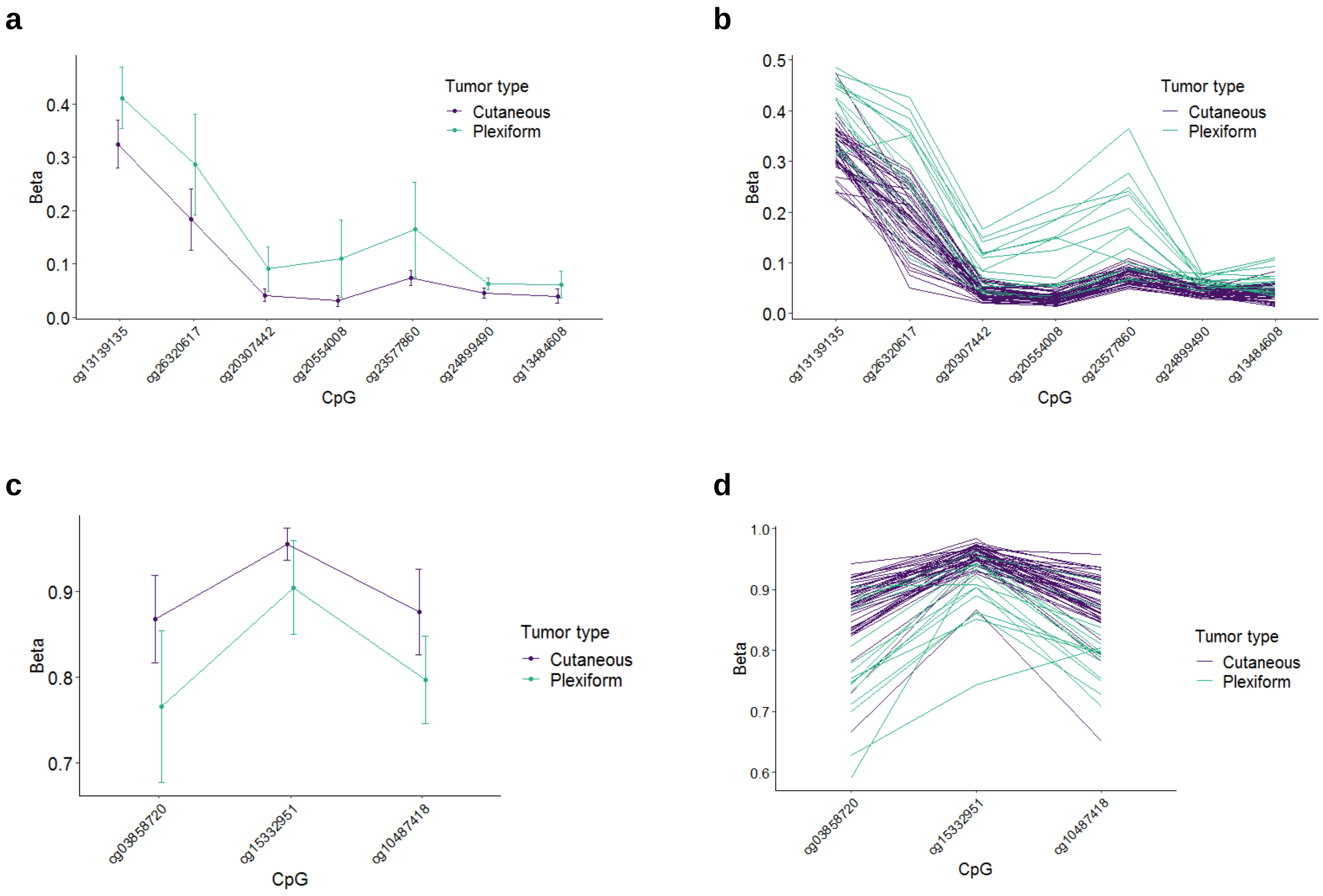


**Supplemental Figure 2. MAP2K3 is differentially methylated between PNFs and CNFs.** a) Mean beta-values at individual positions within MAP2K3 DMR1 plotted by tumor type. Error bars reflect standard deviation. b) Individual beta-values within MAP2K3 DMR1 plotted by tumor type. Each line reflects a single tumor. c) Mean beta-values at individual positions within MAP2K3 DMR2 plotted by tumor type. Error bars reflect standard deviation. d) Individual beta-values within MAP2K3 DMR2 plotted by tumor type. Each line reflects a single tumor.

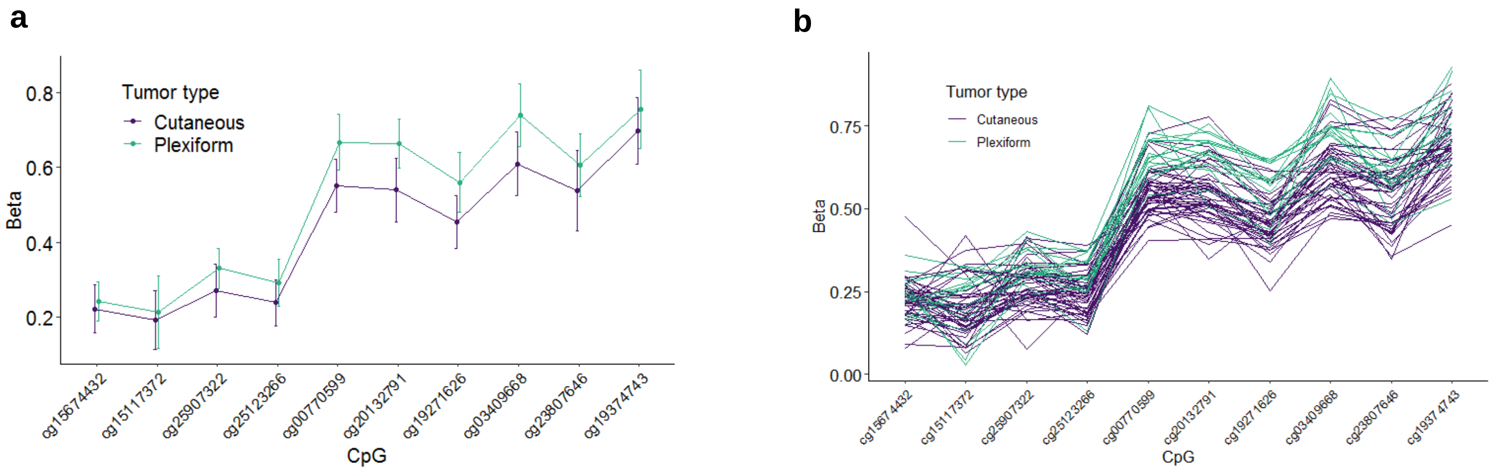


**Supplemental Figure 3. Methylation of MAPK14 is increased in PNFs compared to CNFs.** a) Mean beta-values at individual positions within MAP2K3 DMR1 plotted by tumor type. Error bars reflect standard deviation. b) Individual beta-values within MAP2K3 DMR1 plotted by tumor type. Each line reflects a single tumor.

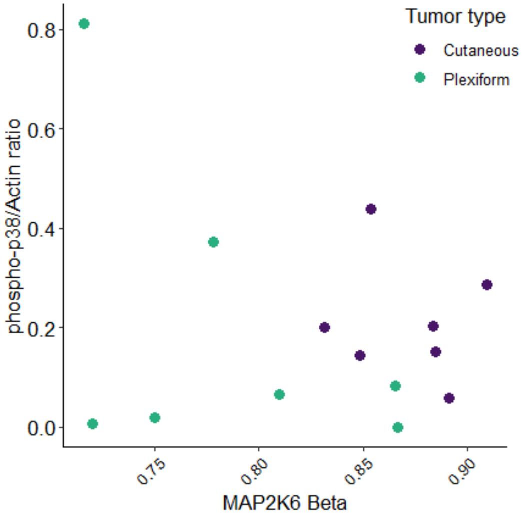


**Supplemental Figure 4. MKK6 methylation does not impact P38 activation.**
Mean beta-values assigned to MAP2K6 (MKK6) for CNFs and PNFs are compared to phospho-p38/actin ratios.
